# Detecting patterns of accessory genome coevolution in bacterial species using data from thousands of bacterial genomes

**DOI:** 10.1101/2022.03.14.484367

**Authors:** Rohan S Mehta, Robert A Petit, Timothy D Read, Daniel B Weissman

## Abstract

Bacterial genomes exhibit widespread horizontal gene transfer, resulting in highly variable genome content that complicates the inference of genetic interactions. In this study, we develop a method for detecting coevolving genes from large datasets of bacterial genomes that we call a “coevolution score”. The method is based on pairwise comparisons of closely related individuals, analogous to a pedigree study in eukaryotic populations. This approach avoids the need for an accurate phylogenetic tree and allows very large datasets to be analyzed for signatures of recent coevolution. We apply our method to all of the more than 3 million pairs of genes from the entire annotated *Staphylococcus aureus* accessory genome of 2,756 annotated genes using a database of over 40,000 whole genomes. We find many pairs of genes that that appear to be gained or lost in a coordinated manner, as well as pairs where the gain of one gene is associated with the loss of the other. These pairs form networks of dozens of rapidly coevolving genes, primarily consisting of genes involved in metal resistance, virulence, mechanisms of horizontal gene transfer, and antibiotic resistance, particularly the SCC*mec* complex. Our results reflect the fact that the evolution of many bacterial pathogens since the middle of the twentieth century has largely been driven by antibiotic resistance gene gain, and in the case of *S. aureus* the SCC*mec* complex is the most prominent of these elements driving the evolution of resistance. The frequent coincidence of these gene gain or loss events suggests that *S. aureus* switch between antibiotic-resistant niches and antibiotic-susceptible ones. While we focus on gene gain and loss, our method can also detect genes which tend to acquire substitutions in tandem or, in datasets that include phenotypic information, genotype-phenotype or phenotype-phenotype coevolution.

## Introduction

Interactions between genes are a major part of evolution, but they are fundamentally difficult to study due to the combinatorial explosion of the number of possible interactions [Phillips, 2008, Mackay, 2014]. In bacteria, widespread horizontal gene transfer creates a much wider range of potential genetic backgrounds and genetic interactions [Arnold et al., 2018]. Detecting gene-gene interactions without performing large numbers of assays requires the development of computational techniques that can handle the necessary volume of genomic data to find signatures in natural genetic diversity.

Methods for finding interactions at the level of genes generally perform Genome-Wide Association Studies (or GWAS) to detect relationships between genes and phenotypes. This approach has been widely used in human populations, and while there have been successes (the first of which was Klein et al. [2005]; see Welter et al. [2014]), GWAS inference in humans is often complicated by the existence of population structure—systematic differences in allele frequencies among subgroups in a population [e.g. Pritchard and Donnelly, 2001, Barton et al., 2019]. This is even more of a problem in bacterial populations, which often have stronger population structure due to their limited and biased recombination [Read and Massey, 2014, Chen and Shapiro, 2015, Power et al., 2017].

There are several existing approaches to detect genotype-phenotype associations in bacteria, the earliest of which are reviewed in Read and Massey [2014]. The software PLINK [Purcell et al., 2007], which is frequently used in human GWAS studies, has also been applied to bacterial datasets [Chewapreecha et al., 2014, Laabei et al., 2014, Power et al., 2016]. Approaches developed specifically for bacteria include those based on regression [Lees et al., 2016, Earle et al., 2016, Lees et al., 2018, Saber and Shapiro, 2020] and those based on phylogenetic convergence [Chen and Shapiro, 2015]. Techniques that explicitly take phylogenetic information into account fare better in highly clonal bacterial systems [Earle et al., 2016, Saber and Shapiro, 2020].

Methods that use phylogenetic convergence are based on homoplasic events on a phylogeny. The package hogwash [Saund and Snitkin, 2020] implements two methods based on ancestral state reconstruction: phyC (introduced by Farhat et al. [2013]) and a more stringent method that was introduced by Hall [2014]. The package treeWAS [Collins and Didelot, 2018] pairs ancestral state reconstruction with simulation given a homoplasy distribution to compute three different tests of association: one that is only uses leaf data and is equivalent to the method proposed by Sheppard et al. [2013], one that is equivalent to phyC [Farhat et al., 2013], and one that is novel and takes into account co-occurance times along the tree. Finally, Scoary [Brynildsrud et al., 2016], uses the method of pairwise comparisons [Maddison, 2000] to find the minimum number of necessary independent co-emergences of two genes given a phylogeny and evaluates association based on this number. These methods are generally computationally demanding, and indeed were left out of a recent simulation study comparing various bacterial GWAS techniques precisely for this reason [Saber and Shapiro, 2020].

While in principle all current published GWAS-style methods could be used to broadly detect gene-gene interactions (by treating the presence or absence of a gene as a “phenotype”), they are in general not built for comparing multiple sets of genes against each other simultaneously and running them for pairwise comparisons of large numbers of genes becomes prohibitively slow. (For instance, it would take treeWAS about 1,200 hours to run on a dataset of the size we consider if split the gene pairs into 5 batches.) Another approach is to specifically design methods for detecting interactions between genes via co-occurrence. Pantagruel [Lassalle et al., 2019] estimates gene trees and evaluates the co-incidence of events on gene trees under a species tree. CoPAP [Cohen et al., 2012, 2013] simulates gain and loss events for pairs of genes along a phylogeny under various coevolutionary models. Liu et al. [2018] use a maximum likelihood method developed by Pagel [1994] to identify genes that have related gain and loss patterns. Most of these approaches use specified evolutionary models, which can become unwieldy over large datasets as tree size grows. The recent method Coinfinder [Whelan et al., 2020] avoids using a full phylogenetic simulation or likelihood analysis by computing the existing phylogenetic statistic of lineage independence D [Fritz and Purvis, 2010] along with a simple statistic of co-incidence to determine putative gene-gene interactions.

Here, we introduce a new method for finding associations between genes in bacterial populations, specifically tailored to accommodate datasets with greater than 1,000 samples, by sidestepping a full phylogenetic analysis entirely. This method, which we call DeCoTUR (Detecting Coevolving Traits Using Relatives), is based on the idea that the clearest signal of biological association is that closely related individuals will differ in their gene presence-absence states in the same way. In our approach, we first identify pairs of closely related individuals. We then find pairs of genes for which, when one gene is gained or lost between a pair of closely related individuals, the other gene is frequently gained or lost as well. We apply our method to the Staphopia database [Petit III and Read, 2018]—which contains over 40,000 publicly available *Staphylococcus aureus* genomes—to detect correlated gain and loss between pairs of accessory genes. The number of such coincident gain/loss events determines a gene pair’s “coevolution score”. We test for interactions by comparing this coevolution score to what would be expected if the two genes were gained and lost independently. With this method, we find interactions between genes involved in a wide variety of functions, including antibiotic resistance, virulence, pathogenicity, phage interactions, mobile genetic elements, and others. The majority of these interactions are positive associations, i.e., pairs of genes that are gained and lost together, rather than substituting for each other. The bias towards positive interactions as well as many of the specific interacting pairs are consistent across genetic backgrounds. We find many interactions between closely linked genes that are likely co-transferred, particularly among genes related to antibiotic resistance. We also find interactions between genes that are not closely linked, especially among genes related to virulence. The coevolution of these pairs is likely to involve multiple transfer events and be driven by epistasis or correlated selection across environments. Finally, we introduce the R package decotur that allows the computation of our coevolution score.

## Methods

### Data

We downloaded all public samples from the Staphopia database [Petit III and Read, 2018], for a total of 42,949 samples. We used the core genome of shared genes determined by Petit III and Read [2018] to compute nucleotide divergences between the samples and we removed 10,308 samples that were identical in core genome sequence and accessory genome composition to at least one other sample. We used each sample’s multi-locus sequence type (MLST, provided by Staphopia) and the publicly-available pubMLST database (https://pubmlst.org/saureus/) to determine its clonal complex (CC). We then performed all subsequent gene-interaction analyses on each clonal complex separately to study the effect of different backgrounds on associations, as well as a combined analysis using a subset of samples from all clonal complexes (see Appendix I for details). We also computed coevolution scores among antibiotic resistance phenotypes across the whole database obtained from ARIBA predictions [Hunt et al., 2017] in Staphopia.

Of the 42,949 public samples in Staphopia, 612 had a sequence type of 0 and were unable to be mapped to a clonal complex. These sequences were added into a “Other” category, along with all sequences that had a known sequence type but no assigned clonal complex. See Figure S1 for sample sizes for each clonal complex.

### Finding close pairs of individuals

We determined closely-related (i.e. “close”) pairs of samples based on the distribution of distances in a pairwise distance matrix—computed using Hamming distances on the concatenated core genome—of all considered samples. This procedure requires a choice of distance cutoff, with pairs of samples whose pairwise distance is below this cutoff are considered to be “close”. In principle, this cutoff can be tuned to whatever scale is of interest, or to match the number of close pairs to the available computational power. We chose cutoffs that resulted in (approximately) 5,000 close pairs for each clonal complex (Table S1) after a preliminary analysis that demonstrated that this number was within a range that yielded relatively consistent results across different cutoffs (Figure S5).

### Filtering genes

For each analysis, we only include genes which have at least two of the less frequent state (presence or absence) in the set of samples used in close pairs. These are the only genes with sufficient presence-absence polymorphism to potentially show a signal of coevolution.

Additionally, previous work has found that the splitting up of gene families dilutes the signal of genetic association with antibiotic resistance phentoypes [Wheeler et al., 2019], and we attempted to mitigate this problem by considering gene “presence” to be the presence of at least one annotation with a particular gene name. For example, in CC1, the gene *mecA* is present in 730 samples out of 1995. In 729 out of those 730 samples, it is present in a single copy, but one sample has two copies. For the purposes of this analysis, we treat those two copies as a single “presence” of *mecA* for that sample.

### Computing the coevolution score

Here we will outline how we test for coevolution between a specific pair of genes, gene 1 and gene 2. To compute the coevolution score, we test each pair of closely related individuals *i* and *j* for evidence of coevolution. Most pairs of close relatives will necessarily be uninformative: for each gene, they will either both have the gene or both lack it, simply by virtue of being closely related. But for genes that are frequently gained and lost, there will be some pairs of close relatives that differ in the focal genes, and these are the pairs that can contribute to the score. Let *P*_*n,k*_ be an indicator variable for the presence of gene *n* in individual *k*, e.g., *P*_1,*i*_ = 1 if individual *i* has gene 1 and 0 otherwise. If one individual has both genes and the other individual has neither, i.e., (*P*_1,*i*_, *P*_1,*j*_, *P*_2,*i*_, *P*_2,*j*_) = (1, 0, 1, 0) or (0, 1, 0, 1), then we add +1 to the score representing a positive association between the genes. Conversely, if one individual has only one gene and the other individual has only the other, i.e., (*P*_1,*i*_, *P*_1,*j*_, *P*_2,*i*_, *P*_2,*j*_) = (1, 0, 0, 1) or (0, 1, 1, 0), then we add +1 to the score representing a negative association between the genes. We compute two separate scores, one for each of these two types of associations. Figure 1 provides an example situation which illustrates how the score focuses on recent co-incident evolutionary events (represented by the red samples in Figure 1), while omitting older evolutionary events.

**Figure 1:**
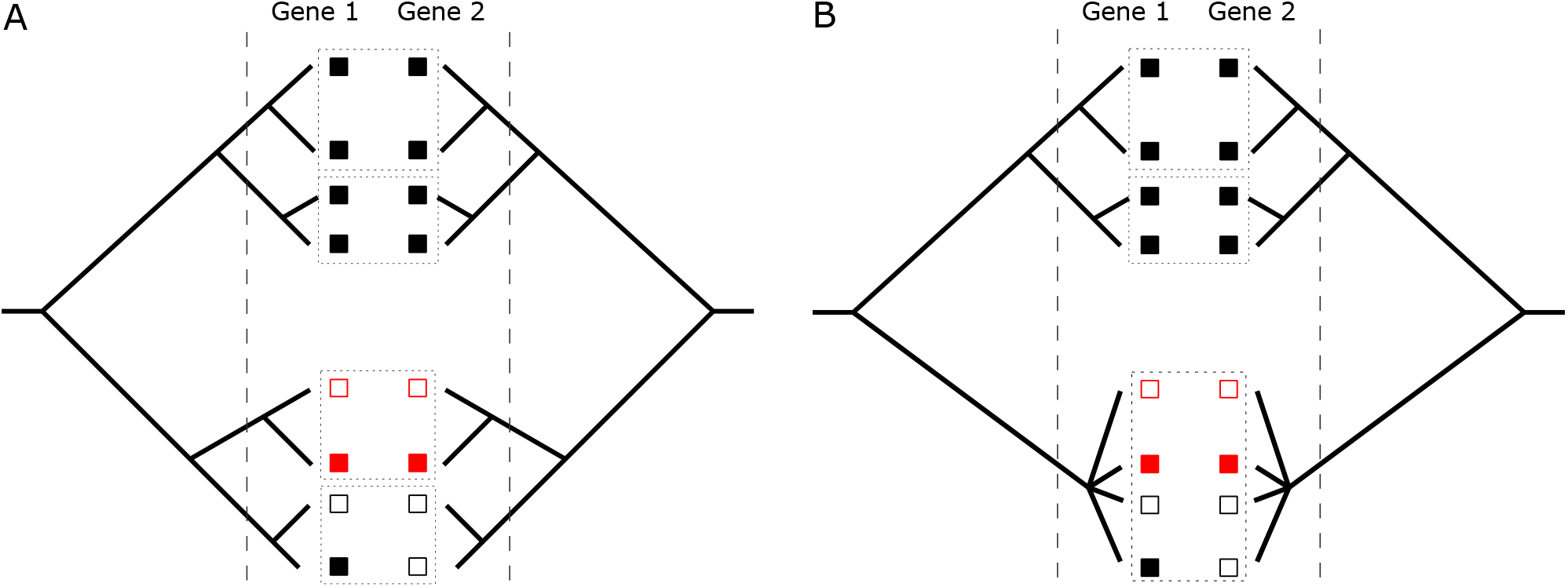
Two examples of the coevolution score computation for a pair of genes (left and right trees in each panel). (A) an example with all disjoint close pairs. (B) An example with an unresolved polytomy “bush,” in which all individuals present are close pairs with each other. The vertical dashed lines indicate the distance cutoffs used to determine close pairs. Filled squares indicate presence of a gene, empty squares indicate its absence. Dashed boxes indicate individuals that are in close pairs with each other. In both (A) and (B), there is exactly one close pair of individuals (in red) that is polymorphic for both genes, indicating recent gain/loss, so only that close pair contributes to the score. The genes differ in the same way (the top red individual has neither gene, the bottom red individual has both), so this contributes to the *positive* score for the gene pair. In (A), the single close pair contributes a value of 1 to the positive score. In (B), this close pair is part of a bush of 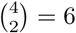 close pairs, so it contributes only 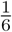 to the positive score. The more ancient event that produced the difference between the top clade (where both genes are present in all individuals) and the bottom clade (where both genes are mostly absent) does not contribute to the score. Note that our method does not actually use the trees, only which pairs of individuals are closely related.

The phylogenies of clonal complexes in *S. aureus* often feature multiple clusters of extremely closely related individuals that form “bushes” in which it is difficult to tell which samples are most closely related (see Figure S4), and for which the specific tree structure may not be difficult to infer accurately. Rather than trying to resolve these bushes, we adjust the value of the contribution for each close pair based on the size of the bush it comes from. Specifically, we partition all the samples into groups where two samples are in the same group if they form a close pair. If pair *k* is in a group with *n*_*k*_ total pairs, then we divide the contribution of that pair to the score by *n*_*k*_. In other words, the maximum total contribution of each bush to the score is 1. This is a very conservative estimate of the amount of coevolution in bushes; it treats a bush as if it were an unresolved polytomy and ignores any tree structure inside the bush that may otherwise indicate a coevolutionary signal. In Figure 1B, there are three bushes, two of size 2 and one of size 4. Only the size 4 bush contributes to the score, and the contribution to the score of that bush is 1/6, as one of the six close pairs in that bush (the red pair) contains a pattern that contributes to the score. Contrast this to the situation in Figure 1A, in which the only bush that contributes to the score is of size 2, so its contribution is 1.

Because our method is based on genetic diversity, it necessarily has the most power to detect coevolution among genes that are at intermediate frequencies. But because we focus on recent/ongoing evolution, the power to detect coevolution does not just depend only on the frequency of a gene in the sample, but also on its distribution. For genes that are essentially exclusively clonally inherited and whose polymorphism corresponds to a deep split in the phylogeny, we do not expect to find a signal, while we have the most power to detect coevolution among genes that are frequently lost or gained via horizontal gene transfer and widely distributed among clades.

Genes that are frequently gained and lost can purely by chance generate a nonzero score. To test for this, we found the total number of discordances between close pairs for each gene. We then used Fisher’s Exact Test to determine if polarized discordances (i.e. discordances that contribute to the positive or negative score) are enriched in any given pair of genes. Pairs of genes that meet a Bonferroni-corrected significance cutoff in this test were kept as statistically significant. Around 33% of our nonzero scores for each clonal complex (166,443 out of 497,772) had a Bonferroni-corrected *p*-value < 0.05. See Appendix C for details. This approach is somewhat liberal, as for a given distance cutoff, some close pairs of individuals will have a genetic distance close to the cutoff and therefore be more likely just by chance to differ at both genes than pairs of individuals that are much closer. In practice, however, we use very tight distance cutoffs so that there is limited variation in genetic distances among close pairs, and we expect this effect to be minor. In datasets with more variation in genetic distance among close pairs, one could use a slightly more sophisticated approach by calculating the rates of gene gain and loss relative to the core mutation rate and use that to determine statistical significance.

To construct interaction networks such as those in Figures 2 and 6, we chose a coevolution score threshold; if two genes have a score above this threshold, we drew a link between them with the weight being the score. These score thresholds were chosen primarily for visualization purposes, but they were always chosen from the extreme high end of the score distribution.

**Figure 2:**
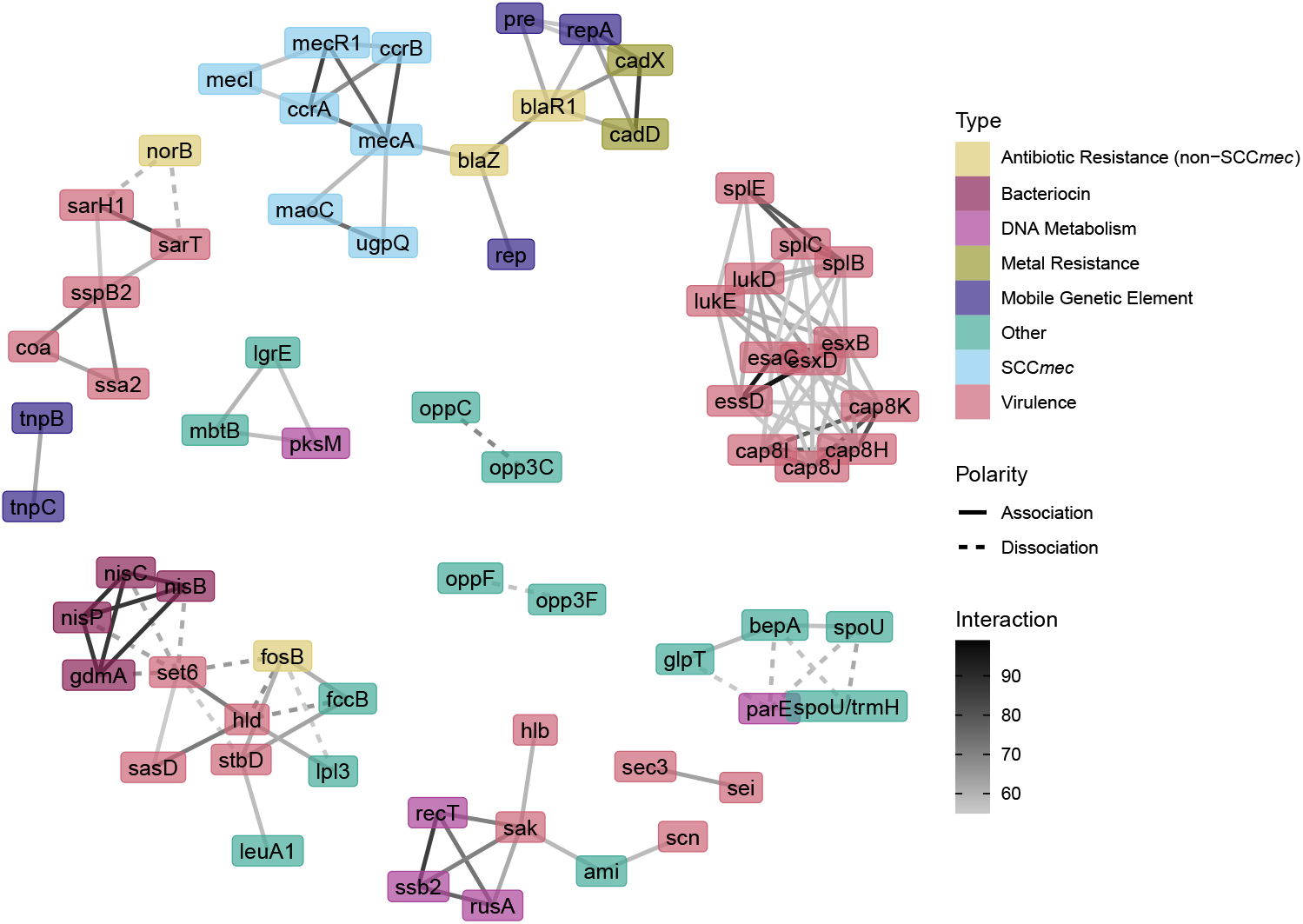
Gene-gene coevolution network for the top 65 significant gene pairs in the full dataset, with nodes colored by gene function, edge color indicating the strength of the inferred interaction, and edge type indicating the polarity of the interaction. A small handful of kinds of genes that are all frequently horizontally transferred—primarily relating to resistance, virulence, or gene transfer itself—tend to dominate the interaction network.

**Figure 3:**
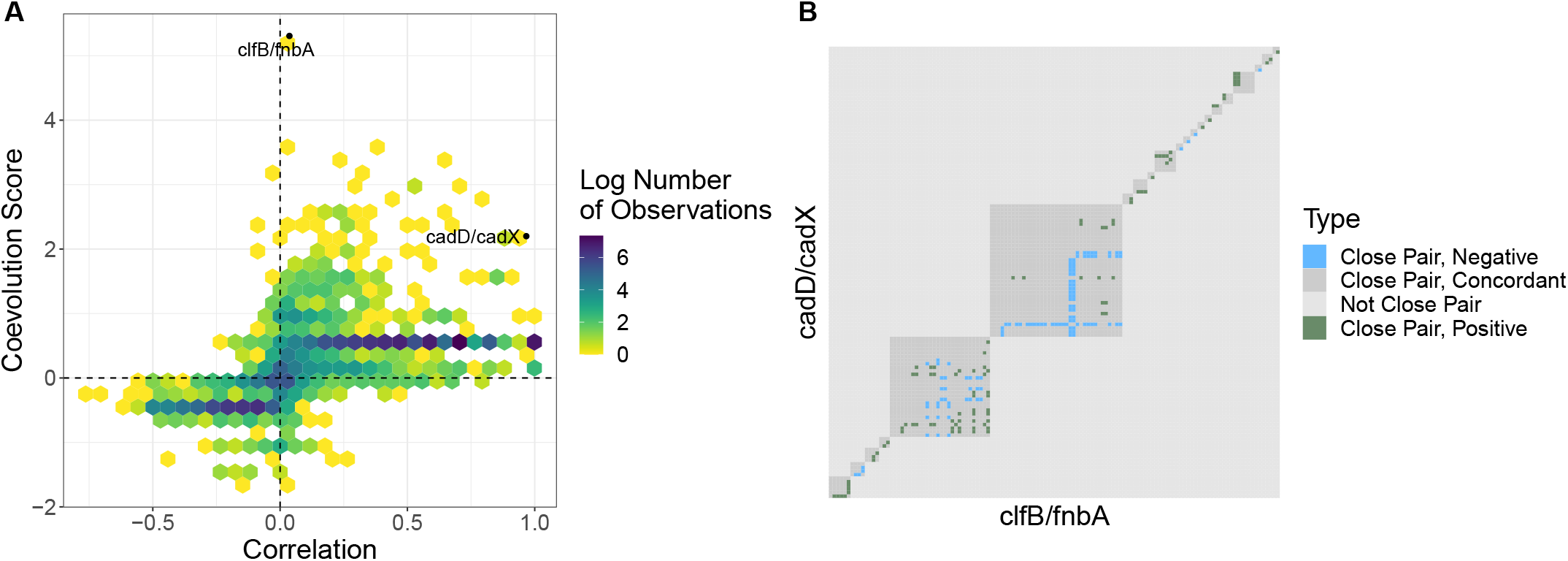
(A) Coevolution scores vs. correlation for CC15. The sign of the score indicates its polarity. The color of a hexagonal bin represents the log of the number of data points in that bin. The coevolution score highlights only a small fraction of the highly correlated pairs of genes, as well as some pairs that do not have a high overall correlation. (B) Interactions between a pair of genes come from multiple sample groups, and different interactions between pairs of genes come from different sample groups. Each x-y coordinate of this heatmap is a sample pair in CC15. The color of each coordinate corresponds to whether that pair contributes to a positive association, a negative association, or not at all (either because it is not a close pair or because it is a close pair but there is no discordance in gene presence/absence). The top part of the matrix corresponds to the *cadD* /*cadX* gene pair, and the bottom part corresponds to the *clfB* /*fnbA* gene pair (labeled in (A)). For both gene pairs, multiple separate groups of closely related individuals contribute to the coevolution score. Different pairs of individuals contribute to the coevolution scores of the two gene pairs.

**Figure 4:**
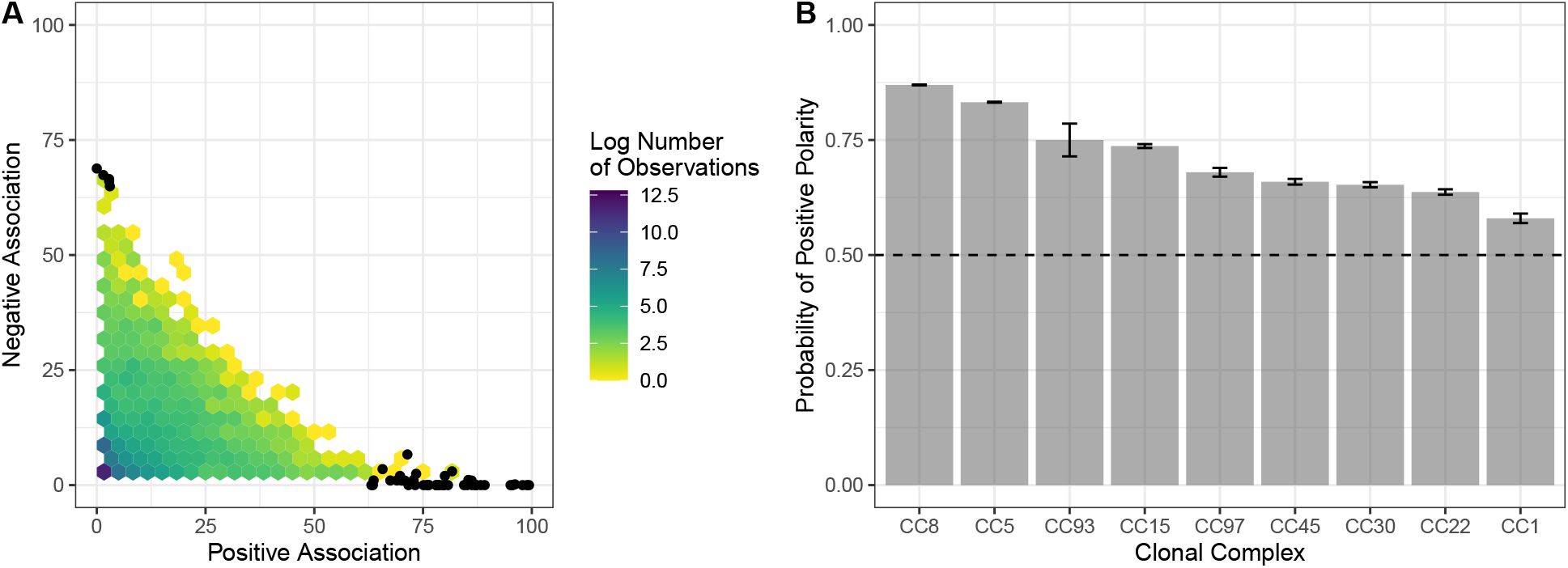
The majority of interactions between coevolving genes are positive. (A) Distribution of positive and negative associations for each pair of genes in the full dataset analysis. Black dots are the top 50 scores. (B) Bars show the estimated probability that the interaction between a pair of genes that are strongly coevolving in the listed clonal complex is positive, i.e., that gain (loss) of one of the genes in the pair is positively associated with gain (loss) of the other gene (see Methods and Appendix E for details). Error bars are the standard error of the probability estimate.

**Figure 5:**
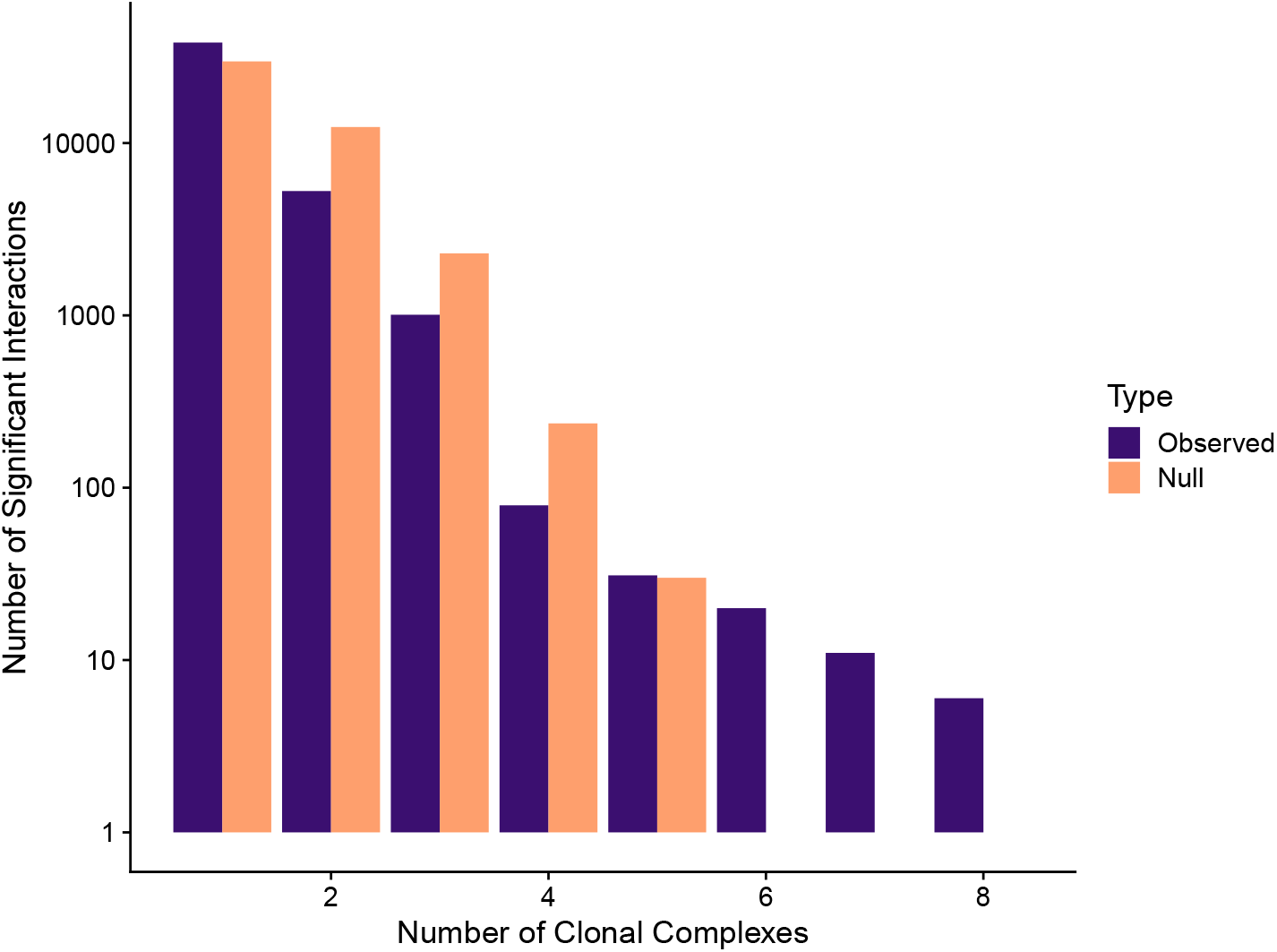
There are a substantial number of significant interactions that appear in the majority clonal complexes at the 95th percentile or higher. This figure plots, per interaction, how many clonal complexes that interaction is above the 95th percentile in score for that clonal complex against a null distribution that is binomial with success rate 5%.

**Figure 6:**
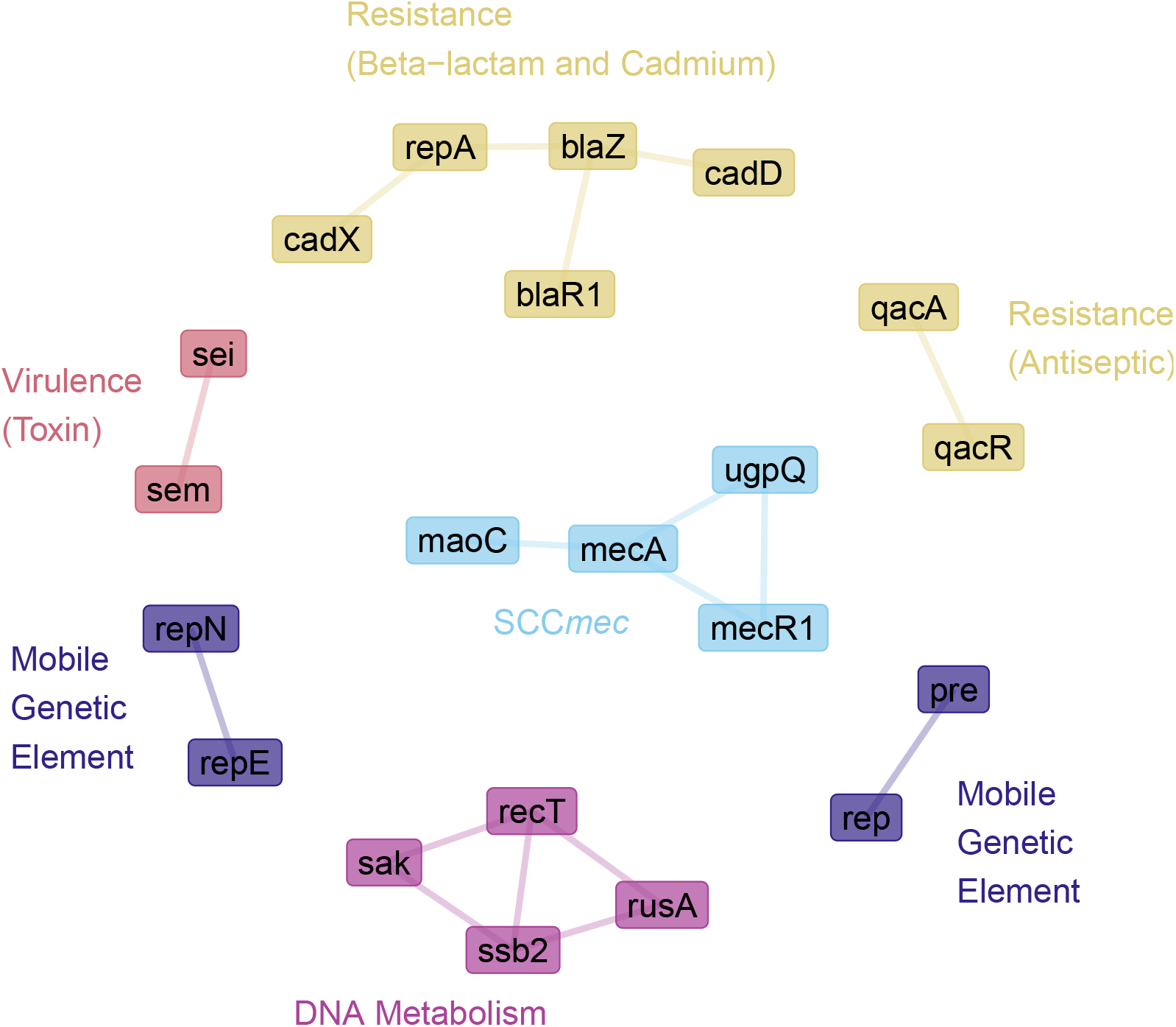
All significant interactions that occur in the top 5% for at least 7 clonal complexes. The groups are labeled and colored by the type of gene they contain. Each interaction in the network has positive polarity; no negative interaction was in the top 5% for more than 3 clonal complexes.

### Detecting positive bias

In the absence of bushes, we have equal power to detect both polarities. But the presence of bushes leads to a bias towards inferring positive interactions (see Appendix E). This bias generally only affects gene pairs with significant contributions from both positive and negative interactions, but to measure the overall distribution of positive and negative interactions—including small ones—we eliminated the bushes by randomly subsampling a single close pair from each bush and computing the score using only those close pairs, and then repeated this process 100 times to achieve 100 independent replicates for the same gene pair. A positive interaction has more positive scores than negative scores across these replicates; a negative interaction has fewer. We then used the resulting distribution of positive and negative interactions to infer the probability of positive polarity in Figure 4 (see Appendix E for details).

## Results

### Gene-gene interactions range from individual operons to complex webs

Throughout all clonal complexes, we consistently find some of the strongest signals of coevolution among genes related to resistance to antibiotics and metals; mobile genetic elements; and genes that influence virulence and toxicity, by e.g. producing a toxin, being involved in biofilm formation, or regulation. But the coevolution networks also include many genes whose functions do not obviously pertain to any of the aforementioned functions. Figure 2 provides an example of such an interaction network obtained from a full-dataset analysis, using only interactions in approximately the top 0.01% of scores that passed the significance test outlined in Appendix E. The procedure for obtaining this full-dataset analysis is outlined in Appendix I.

There are five notable large clusters of interactions in Figure 2. The largest contains all of the major genes contained in the SCC*mec* cassette, a non-SCC*mec* operon that also confers beta-lactam resistance (*blaZ* and *blaR1*), a cadmium resistance operon (*cadD* and *cadX*)—reflecting a known plasmid interaction [McCarthy and Lindsay, 2012]—and genes that are involved in plasmid replication (*pre, rep*, and *repA*). The next largest is a collection of virulence genes, including toxin-producing genes (*lukE, lukD, essD, esxD, esaC*, and *esxB*), endopeptidases that are regulated by *arg* (*splB/C/E*), and capsule genes (*cap8H/I/J/K*). There are two other virulence-based clusters, one of which has a negative interaction with *norB*, which confers quinolone resistance. The other virulence-based cluster also contains genes involved in DNA metabolism (*recT, rusA*, and *ssb2*). Finally, the large major cluster contains virulence genes, antibiotic resistance genes, and bacteriocin genes (specifically, the lantibiotic nisin). There are also six smaller clusters of genes of varying function. These interactions paint a picture of recent genetic coevolution in *S. aureus* that focuses on host-pathogen interaction in all of its many facets.

We also note that a handful of interactions seen in Figure 2 are artifacts of annotation. In particular, *spoU* and *trmH* are two names for the same gene. In addition, *opp3C* and *opp3F* refer to specific alleles of *oppC* and *oppF*. The “negative” interactions between *spoU* and *spoU/trmH, opp3C* and *oppC*, and *opp3F* and *oppF* are all due to the fact that some studies use one name and some studies use another. This inconsistency is a challenge for any large-scale analysis of genomic content; fortunately it is frequently easy to spot in results like these.

### Coevolution score differs substantially from correlation

An easy-to-compute first pass at attempting to detect genetic interactions is to compute the correlations between presence-absence vectors for pairs of genes, without performing any phylogenetic correction. Figure 3 compares our coevolution score with this correlation for each pair of genes for samples in CC15, and Figure S3 shows this comparison for each clonal complex.

Overall, there is an association between the two measures, as is to be expected: all gene presence-absence configurations that contribute to the coevolution score also contribute to correlation. But the converse is not true, and indeed, most highly correlated pairs of genes have modest coevolution scores; in other words, most correlation appears to be phylogenetic. Thus, coevolution score can be used to filter out the vast numbers of highly correlated gene pairs to focus on the few currently or recently coevolving ones. There are few gene pairs that have high coevolution score but low correlation. This is because the coevolution score is driven by exceptional events (double gene gains or losses between extremely closely related individuals). Even a handful of such events can provide a clear signature of coevolution, while being too rare to produce a strong correlation. The fact that these high-score, low-correlation pairs of genes are rare suggests that ancient, long-term evolution is concordant with recent, short-term evolution.

Two high scores in CC15 in Figure 3A are between *cadD*-*cadX*, two genes involved in cadmium resistance that are found on SCC*mec*, and between *clfB*-*fnbA*, two genes that are involved in cell surface adherence and host colonization. We chose these two pairs to compare because they represent a high-score, high-correlation pair (*cadD/cadX*) and a high-score, low-correlation pair (*clfB/fnbA*). Figure 3B shows all samples that contributed to the scores for each of these pairs. The samples in CC15 are the rows and columns of the matrix, and each square represents a pair of samples. Close pairs of samples are shown in the darker gray squares and are colored by their contribution to the score. For each of these two pairs of genes, contributions come from multiple different groups of close pairs, and these groups contribute different amounts for the different interactions.

### Most associations between genes are positive

Our method can detect both positive coevolution (e.g., where the gain of one gene is associated with the gain of the other) and negative coevolution (e.g., where the gain of one gene is associated with the *loss* of the other). Because we measure both positive and negative interactions for the same pairs of genes separately, we can identify pairs of genes that have strong positive interactions in some parts of the tree and strong negative interactions in other parts of the tree. In general, we would expect this effect to occur more frequently for larger sampling scales (i.e. multiple clonal complexes or a large clonal complex) and less frequently for smaller sampling scales (a single, small clonal complex). Figure 4A shows that in a full-dataset analysis, while the strongest interactions are primarily confined to mostly-positive or mostly-negative, there are some interactions of notable magnitude with contributions from both. Figure 4B displays the probability that an interaction is positive for each clonal complex, correcting for bush-induced positive bias (see Methods and Appendix E). We find that positive interactions are significantly more likely than negative interactions in all clonal complexes (Figure 4).

### Strong interactions are consistent across backgrounds

Each clonal complex reflects a different “path” of evolution for *S. aureus*, potentially facing different environments and different selective pressures. To determine whether the same interactions consistently appear across the clonal complexes, we tabulated the number of times each interaction appeared in the top 5% of scores for each clonal complex. We then compared the distribution of the number of clonal complexes for which each interaction appeared in the top 5% to a null distribution where the top 5% was chosen randomly from the set of interactions for each clonal complex independently. This null distribution is a binomial distribution with probability 5% conditional on one success.

Figure 5 plots these two distributions. The observed distribution has many more interactions that are strong in > 5 clonal complexes (and fewer that are strong in < 5) than would be expected if the interactions were independent across clonal complexes. Thus, strong interactions are more likely to be strong across clonal complexes, and so these interactions are consistent across clonal complexes.

Figure 6 displays the 28 significant interactions that are in the top 5% of scores in at least 7 clonal complexes. These interactions can be divided into 8 disjoint groups: the SCC*mec* cassette, one biofilmrelated virulence group, two toxin-related virulence groups, one bacteriocin group, one antiseptic resistance group, one beta-lactam and cadmium resistance group, and one group with genes that are involved in various DNA activities (replication, recombination, restriction).

### Antibiotic resistance phenotypes fall into two sets of interactions

Our coevolution score is not restricted to gene presence/absence and can be applied to any binary trait. We initially applied the score to SNPs, but found that accessory genes had more interesting evolutionary patterns in this dataset. We can also apply our method to binary phenotypes, such as the presence or absence of antibiotic resistance. Staphopia predicts antibiotic resistance phenotypes using ARIBA [Hunt et al., 2017]. For each sample in the full-dataset analysis, we computed coevolution scores for these predicted antibiotic resistance phenotypes. Figure 7 displays a heatmap of the significant interactions and significant pairwise correlations for these phenotypes. Note the the coevolution scores as scaled so that the highest magnitude is one and the lowest magnitude is zero.

**Figure 7:**
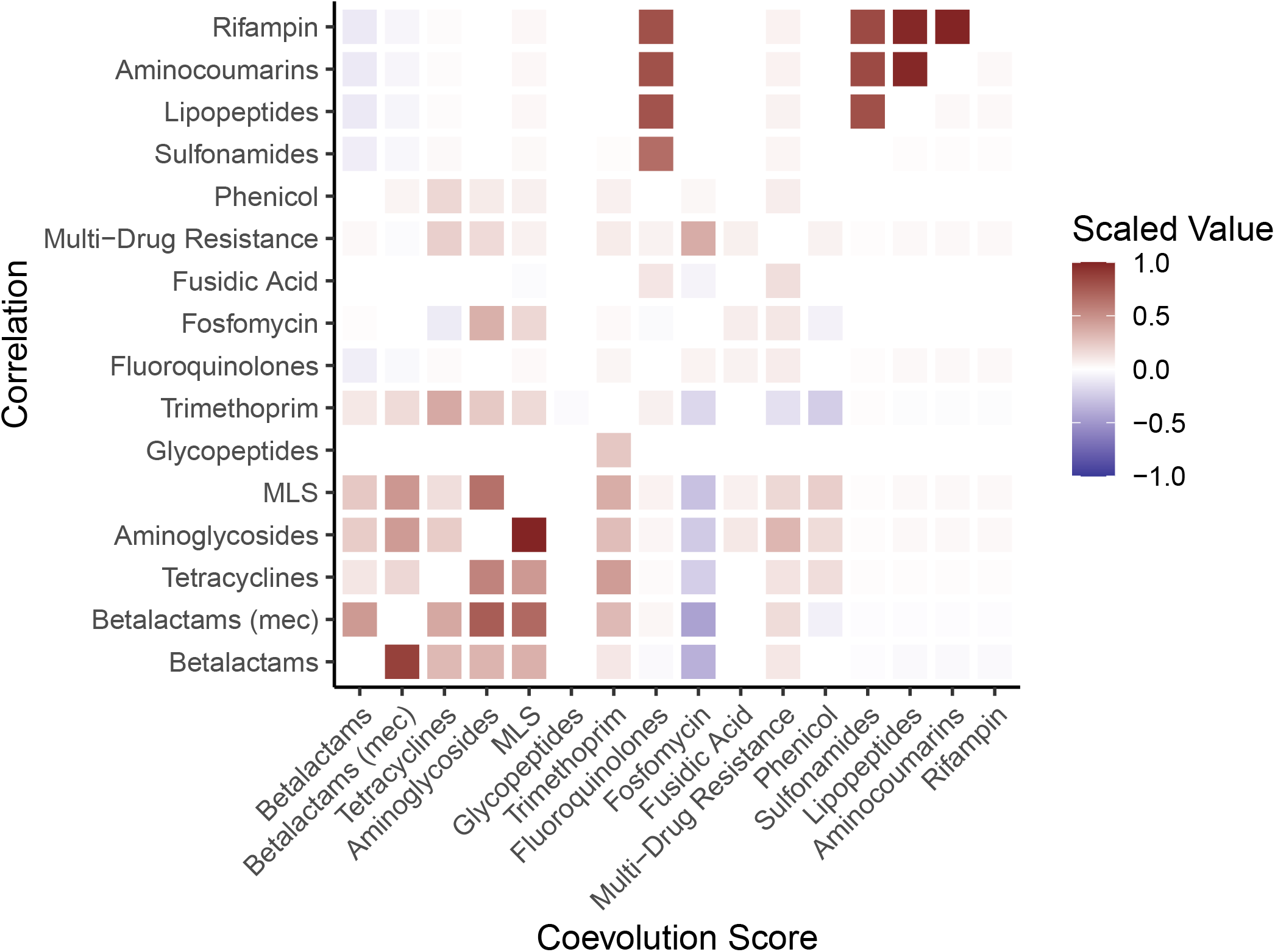
Significant coevolution scores (bottom-right triangle) and pairwise correlation values (top-left triangle) for predicted antibiotic resistance phenotypes in Staphopia. The strongest interaction block involves resistance to MLS, aminoglycosides, betalactams, and tetracyclines. Multi-drug resistance, and resistance to fusidic acid, glycopeptides, trimethoprim, fluoroquinolones, rifampin, lipopeptides, and sulfonamides have peripheral interactions. The axes are ordered according to a hierarchical clustering on the coevolution score.

There is a strong positive interaction cluster between both beta-lactam resistance phenotypes, MLS, aminoglycoside, trimethoprim, tetracycline, and phenicol resistance. The two strongest interactions are between aminoglycoside and MLS resistance and between SCC*mec* and non-SCC*mec* beta-lactam resistance. Fosfomycin resistance appears to strongly negatively interact with the other resistances. Finally, the remaining resistance phenotypes form a peripheral, weakly interacting group. These phenotypes are also in general much rarer than those in the beta-lactam interaction group, so their signal is limited.

The high-scoring group also has high correlation, but fosfomycin resistance has a clear negative signal with the coevolution score and no clear signal with correlation. Five of the peripheral resistance phenotypes are strongly correlated with each other, but have very little signal with the coevolution score.

### Comparison with Coinfinder

To compare our method with that of Coinfinder, we ran Coinfinder on CC97. Figure 8 compares an interaction network produced by Coinfinder with one produced by our method for this clonal complex. One major distinction between our method and that of Coinfinder is that we use the genetic distance cutoff as a way to avoid having to deal with the phylogeny, whereas Coinfinder handles phylogeny-induced correlations by presenting Fritz and Purvis [2010]’s D statistic as metadata for their interaction network. However, the phylogenetic statistic D used by Coinfinder uses some of the same ideas that we use in motivating our score: in particular, situations that maximize D (i.e. all sister taxa differ in gene presence/absence) are necessary but not sufficient for maximizing our score (which also takes coincident difference between genes into account). Our use of the distance cutoff eliminates the major computationally difficult step of Coinfinder—computing D for each gene using the whole tree—at the cost of only looking at recent events.

**Figure 8:**
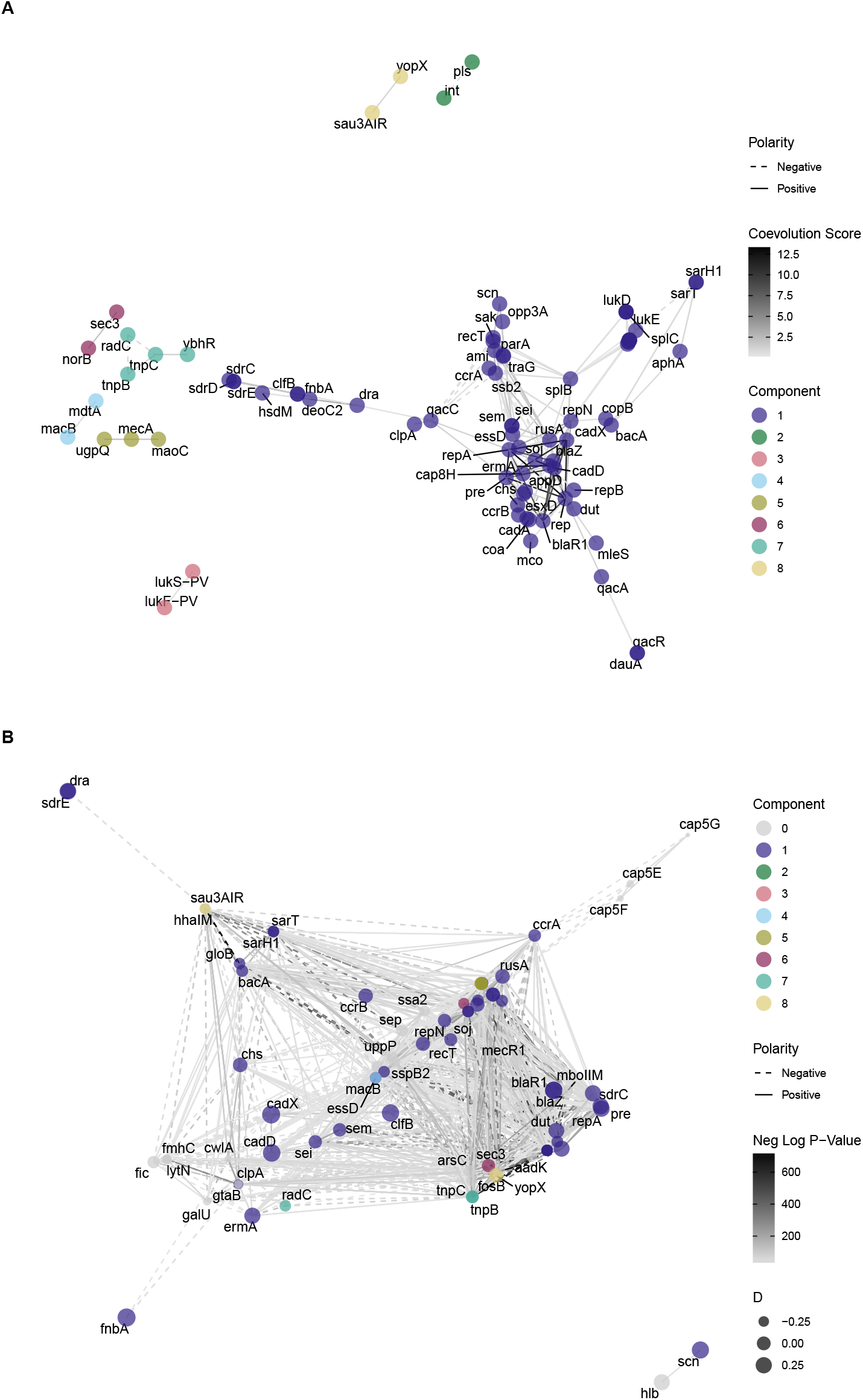
Our results show some overlap but some differences with those of Coinfinder when run on the same clonal complex (CC1). (A) Significant coevolution score network, with score cutoff at the 97.3rd percentile. (B) Coinfinder network, with the p-value of the interaction as a proxy for interaction strength, and the phylogenetic signal score D reported as node size (largest is more phylogenetically independent). Node color corresponds to component in the coevolution-score network (A). Nodes in (B) that are not in (A) are labeled as component 0.

Figure 8 compares an interaction network obtained using our method with Coinfinder’s default method. We used a score cutoff of the 97.3rd percentile to obtain 83 interacting genes, which is close to the 84 interacting genes obtained by Coinfinder. While there are significant overlaps, implying that some signals are detected by both methods, there are also substantial differences in the results of the two methods. In particular, both methods detect many of the same genes as interacting, but the underlying network structure is very different and communities are not preserved across methods. This lack of coherent community structure in Coinfinder may be due to the fact that there is no sense of interaction “weight” (only a p-value), and so Coinfinder reports many more interactions that are potentially weak, which would obscure community structure in the interaction network.

## Discussion

We have presented a new method for detecting interactions between genes in large bacterial datasets, using pairwise divergence in the core genome to find closely-related pairs of organisms and finding pairs of genes that differ within the same close pairs. We applied this method to Staphopia, a dataset of more than 42,000 genomes of *Staphylococcus aureus*, to a find a network of accessory genes that are being gained and lost together.

The gene interactions that our method detects present an interconnected picture of various ways in which *S. aureus* interacts with its environment. Along with antibiotic resistance genes, we found substantial interaction with genes that promote virulence and pathogenicity—ranging from host colonization to toxin production—as well as genes that code for resistance to metals and genes that are involved in plasmid replication, bacteriocins, and DNA metabolism. Our results suggest that recent gene-gene coevolution in *S. aureus* is a complex, interconnected web in which horizontal gene transfer allows lineages to rapidly acquire a suite of traits involved in pathogenicity, including antibiotic resistance, host colonization, and competition with other bacteria.

The gene interactions we detected were frequently, but not universally, conserved across different clonal complexes. The different environments that different clonal complexes have recently encountered may lead to differences in the effects of various genes on other genes through different selection pressures and different pleiotropic effects. Also, differences in horizontal gene transfer between clonal complexes may have led to different opportunities for interaction. Studying the differences in clonal complexes with respect to genetic interactions and horizontal gene transfer may reveal important information about recent *S. aureus* evolution. We found that most interactions between pairs of genes are positive, with the presence of one gene correlated with the presence of the other, rather than anti-correlated. This is similar to the result found by Hall et al. [2021] using a different method (Coinfinder) in a different system (*E. coli*), suggesting that it may be a general pattern. Both of these results support the idea that HGT-based evolution is driven more by the collection of genes that work well together as opposed to the sorting of a diverse set of genes that are interchangeable. Of course, selection may favor linking such sets of genes into operons, which will then facilitate their co-transfer and strengthen the pattern of positive associations.

One of the more unexpected results we found was cadmium resistance’s frequent strong coevolution with antibiotic resistance. It is not obvious why these genes should have such a strong signal across clonal complexes, especially considering that there are other genes that are also frequently found in SCC*mec* that show much weaker interaction. One potential explanation could involve a linkage of cadmium resistance to survival in wastewater as a transmission mechanism [Amirsoleimani et al., 2021, e.g.].

One major question about the mechanism of genetic interaction that is generally difficult to achieve without extremely thorough genome assembly annotation is the distinction between interactions between linked genes—which would imply a single gain/loss event—versus interactions due to unlinked genes, which would imply multiple gain/loss events. Outside of analyzing known operons, this distinction is impossible to properly interrogate in most cases without comparing the specific alignment of each genome simultaneously, which is intractable for large datasets. For unlinked genes, a major question is if there is a consistent order in which the gain/loss events occur within a pair of interacting genes. For instance, among genes that interact with antibiotic resistance genes, we would expect potentiating genes or mutations to tend to be acquired before the resistance gene, and compensatory genes or mutations to be acquired after. While we cannot address this question with our pair-based approach, it may be possible to extend it by using local phylogenies to infer the order of gains and losses.

The Staphopia database is compiled from public data; sampling biases in these data will therefore be preserved in Staphopia. One major such bias is the overabundance of MRSA (methicillin-resistant) vs. MSSA (methicillin-sensitive) strains due to the important clinical relevance of certain MRSA strains. This bias could potentially inflate the importance of the SCC*mec* cassette. Two aspects of our method can mitigate this bias. First, by downweighting bushes by their size, we avoid the score being dominated by a recent well-sampled branch of the tree. Second, by splitting up some of our analyses by clonal complex and then tracking interactions that occur consistently across clonal complexes, we mitigate effects of uneven sampling in clonal complexes. The only true solution to this bias, however, would be to design studies that deeply sampled genomes in a way that somehow reflected the underlying population structure of *S. aureus* and reduced biases away from strains with particular antibiotic resistance and virulence characteristics. The problem with this solution is that we do not know the actual underlying population structure, so perhaps more scattershot metagenomic sampling will provide an alternative set of differently-biased samples for comparison.

A major limitation of compiling and analyzing genomic data from multiple sources is inconsistencies with gene annotation. Potentially incomplete, ambiguous, or mismatched annotation reduces the power of methods like the one presented here to detect interactions, and we see that it can also produce spurious interactions. However, it is worth noting that one of the advantages of a database like Staphopia in the first place is more consistent annotation, with genomes annotated at the same time using the same software or database. In this work, we limited ourselves to only those genes that were annotated in Staphopia by way of being assigned a “name” in the standard sense (like “*mecA*”). This technique is effective at quickly obtaining broad-scale results, but analyses on a finer scale would require additional steps to mitigate this limitation.

The ability to discover and investigate interactions between genes in bacteria will only increase with the increase in the accessibility of large amounts of data provided by databases such as Staphopia. With more data, we may be able to discover more interactions with smaller signals, or interactions that are strong but rare. We constructed our method specifically to be able to keep up with this progress. Methods such as ours, coupled with databases such as Staphopia, will allow both the study of broad-scale patterns of bacterial evolution as well as providing more focused results for future study.

## A Sample sizes for each clonal complex

Figure S1 displays the sample sizes for each clonal complex in this dataset, both prior to and after removal of samples that were identical across both the core genome and in accessory gene presence/absence.

**Figure S1:**
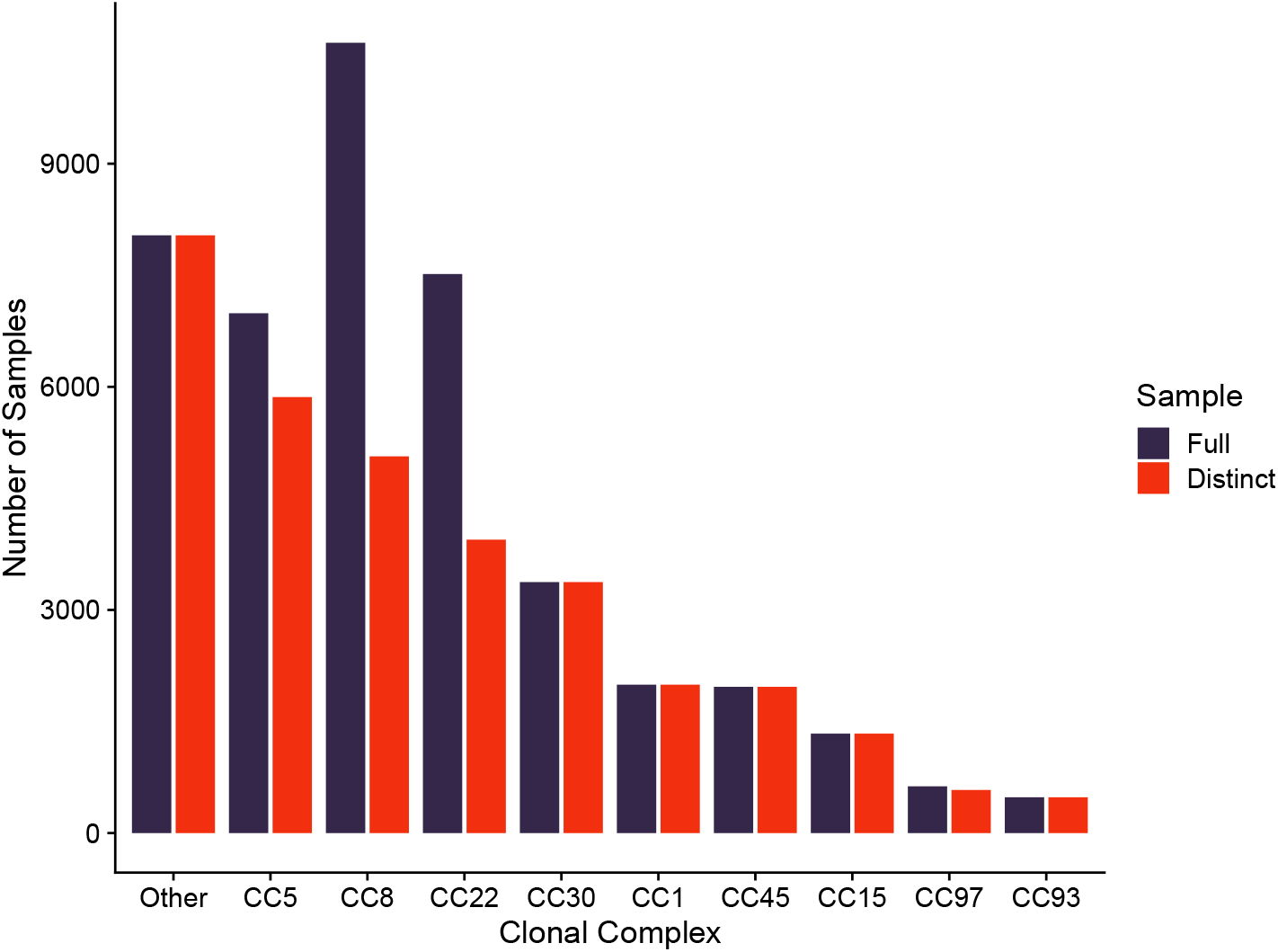
The total number of public samples per clonal complex in Staphopia (“Full”), as well as the total number of samples obtained after removing samples that were identical across both the core genome and accessory gene presence/absence (“Distinct”).

## B Distance scales and number of close pairs for each clonal complex

For our data, the individual clonal complexes varied as to the scale of divergence they comprised. Table S1 displays the distance cutoffs and number of close pairs used in our analyses of individual clonal complexes. Figure S2 displays the pairwise distance distributions for all distinct samples in each clonal complex, along with the distances cutoffs from Table S1.

## C Statistical test for significance

Because the coevolution score only records events that affirmatively contribute, it is possible for pairs of genes that individually vary frequently across the set of samples to accumulate a substantial score by chance. To test for this, for each gene in each clonal complex, we first compute the number of presence/absence discordances that gene has across all close pairs. Then, for each pair of genes, we use Fisher’s Exact Test to see if the number of joint discordances (i.e. if both genes are discordant for the same close pair) is significantly greater than would be expected by chance. We use the most conservative multiple testing correction (the Bonferroni correction) with *α* = 0.05 to obtain significant interactions.

**Table S1:** Core genome distance cutoffs and number of close pairs for each clonal complex. For CC8 and CC22, the close pairs were downsampled to reach the target number of 5000 due to the fact that there were more than 5000 pairs of samples that were identical in the core genome. Distances are fraction of divergent bases in the core genome.

| Clonal Complex | Distance Cutoff | No. Close Pairs |
| --- | --- | --- |
| CC97 | $4.98 \times 10^{-5}$ | 5125 |
| CC93 | $3.84 \times 10^{-5}$ | 4847 |
| CC15 | $2.13 \times 10^{-5}$ | 4368 |
| CC1 | $9.96 \times 10^{-6}$ | 4211 |
| CC45 | $8.54 \times 10^{-6}$ | 4177 |
| CC30 | $1.42 \times 10^{-6}$ | 3566 |
| CC5 | 0 | 4397 |
| CC8 | 0 | 5000 |
| CC22 | 0 | 5000 |

**Figure S2:**
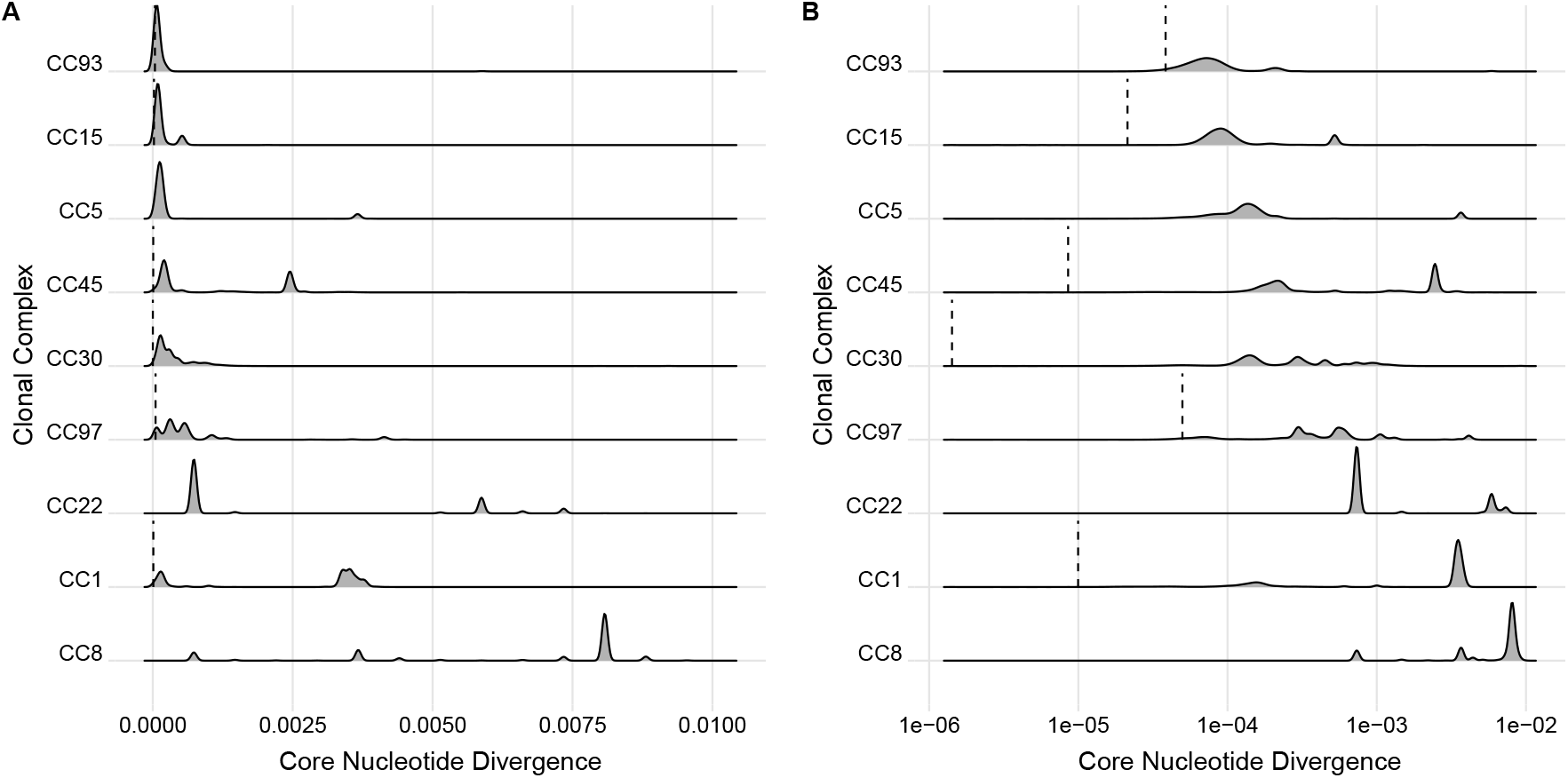
Pairwise nucleotide divergence in the core genome of Staphopia samples, where all zero values were excluded. (A) Linear divergence scale. (B) Log divergence scale. Distance cutoffs used to find close pairs are given by vertical dashed lines.

## D Score vs. correlation for all clonal complexes

Figure S3 displays coevolution score vs. correlation for each clonal complex. The patterns seen in Figure 3 are consistent across the clonal complexes.

**Figure S3:**
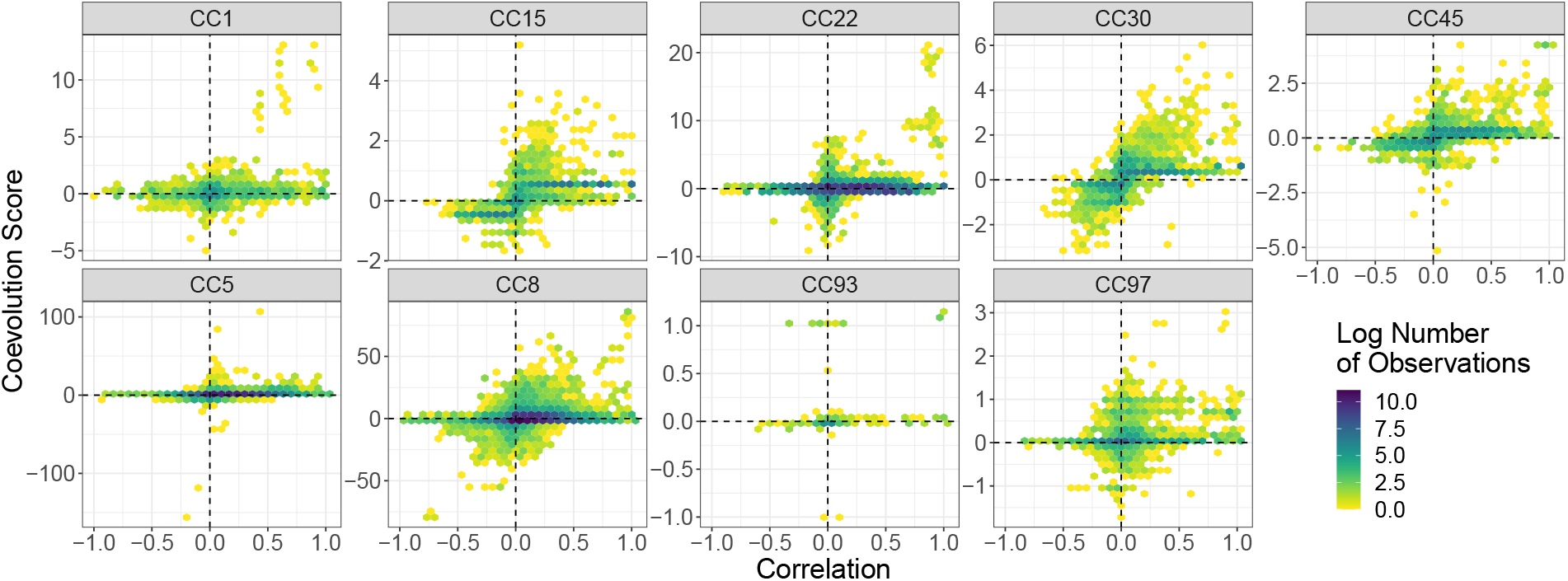
Coevolution score vs. correlation for each clonal complex. The sign of the score indicates its polarity. The color of a hexagonal bin represents the log of the number of data points in that bin. The coevolution score highlights only a small fraction of the highly correlated pairs of genes, as well as some pairs that do not have a high overall correlation.

## E Positive Bias when close pairs are not all disjoint

If close pairs share samples between them, they are no longer independent. This effect is particularly strong in bushes, where each sample is in *n* close pairs, where *n* is the size of the bush. This dependence may affect the balance of positive and negative polarities among the scores. To properly estimate whether or not genetic interactions are biased towards positive association, we must correct for potential positive bias due to this dependence. We correct for this bias by (for the purposes of this portion of the analysis only) re-computing the coevolution score using only disjoint sets of close pairs, so that no close pair shares a sample with another close pair.

To create sets of independent close pairs for the purposes of detecting the prevalence of positive interactions, we randomly chose one representative close pair for each bush, computed the scores across those selected disjoint sets of close pairs, and repeated that process 100 times. We then noted for each gene pair if its positive score was bigger than its negative score more often across these 100 replicates. If so, that interaction was labeled “positive”. We fit a Bernoulli distribution to the distribution of these positive labels. The probability parameter of this distribution represents the probability that a particular interaction is positive.

## F Example phylogenetic tree for CC97

Figure S4 displays the phylogenetic tree for CC1, the clonal complex with the largest sample size in Staphopia. The bushy structure is clearly seen, with the vast majority of samples belonging to a handful of very-recently-diverged bushes.

## G Gene annotation

We used the gene names provided by Staphopia for gene annotation. However, due to the fact that Staphopia is a collection of publicly-available datasets with no consistent curation, it is possible that a particular gene does not have its name category filled out.

For each gene name in Staphopia, we found the corresponding gene product annotations. The gene product is the highest level of annotation for a gene in Staphopia that includes some information about the gene itself (other possible annotations are NCBI locus tag and product ID, both of which are narrower). We then found all instances of the gene product in Staphopia, and tabulated the number of those instances that were unnamed. If there exist unnamed instances of the same gene, and the gene product is known, then these instances would appear in this set. Table S2 lists all the genes that have *any* other instance of their gene product that occurs without a name. There are 144 such genes.

**Figure S4:**
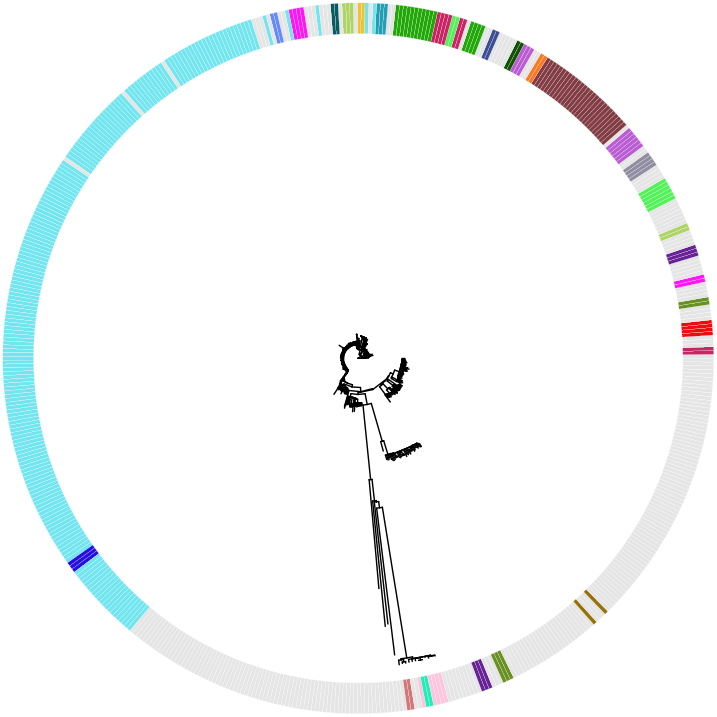
The phylogenetic tree of CC97 consists of bushes of very closely-related samples separated by long branches. This bushy structure makes the specific relationships between samples within a bush subject to resolution error. Colors distinguish separate bushes for close pairs at distance cutoff 5 × 10^*−*5^. Samples not included in a close pair are not colored.

For each gene product in Table S2, we observed the distribution of lengths of all genes corresponding to that product, separated by name. If the distribution of gene lengths with no name clearly matched that of a named gene, we renamed the genes with no name after that name. The genes affected by this were: *add, ahs2, AHS2, alr, apbC, arcB, aspC, blaR1, chrR, coa, hadL, hmrA, hyuA, lytA, map, metN, polA, prfC, radC, sdrD, sdrE, sph, strH, ugl, yidD, ykuR*, and *yoaB*. This adjustment is likely to turn absences into presences, which means genes that may have had substantial positive and negative relationships may find the negative relationships spurious due to lack of annotation.

In addition, if the length distribution of a gene product subsumed that for a named gene, we added that gene product to our list of “genes” in order to make sure we do not miss a gene due to poor annotation. The genes affected by these were: *atsA, chuR, dctP, dld, hysA, int, nhaC, paaK, relA, sotB, tadA, tagE, tetR, traA*, and *ybgI*. These adjustments do not affect results for the named genes, but introduce additional tests of association with the corresponding gene products.

**Table S2:**
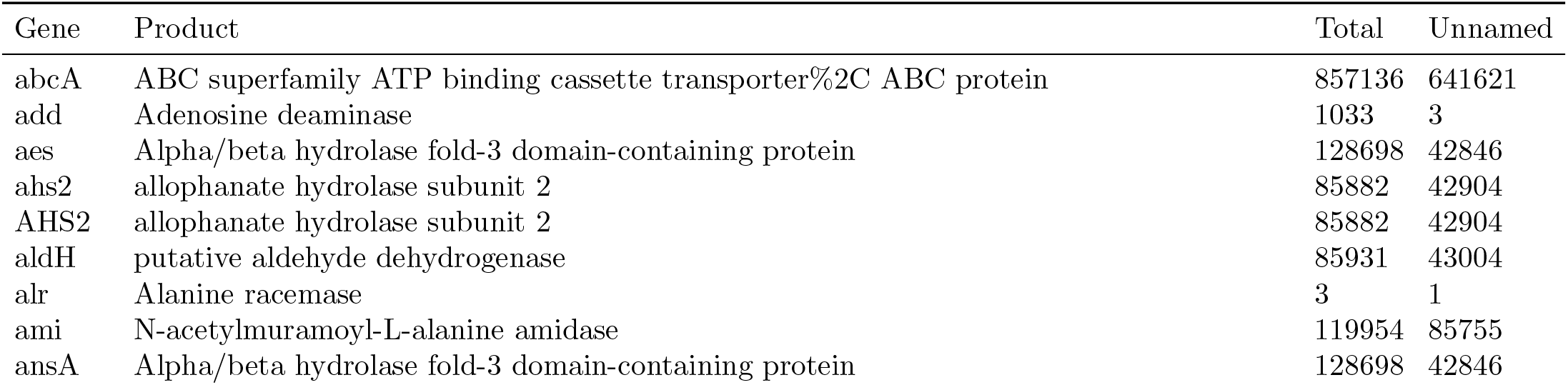
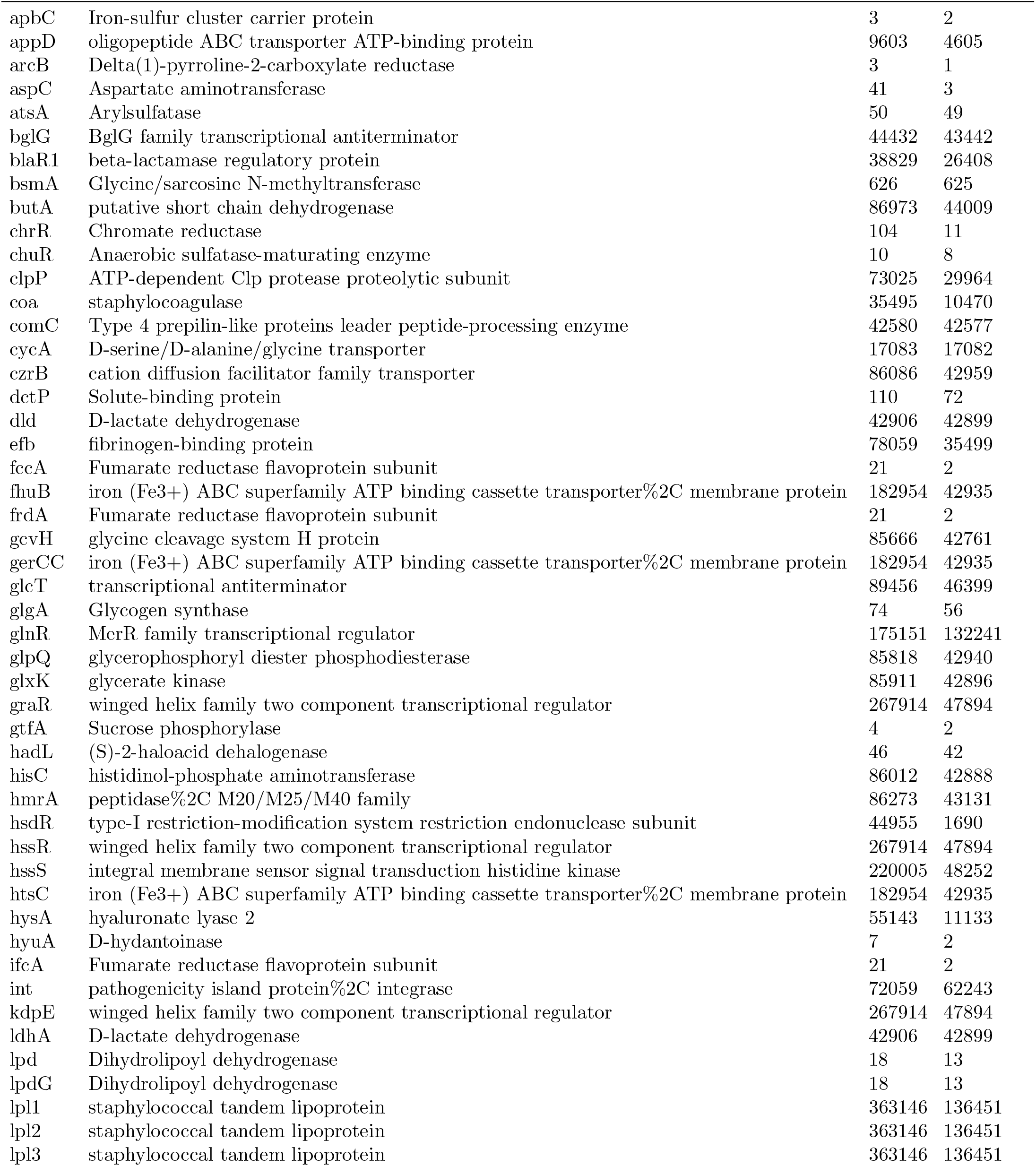
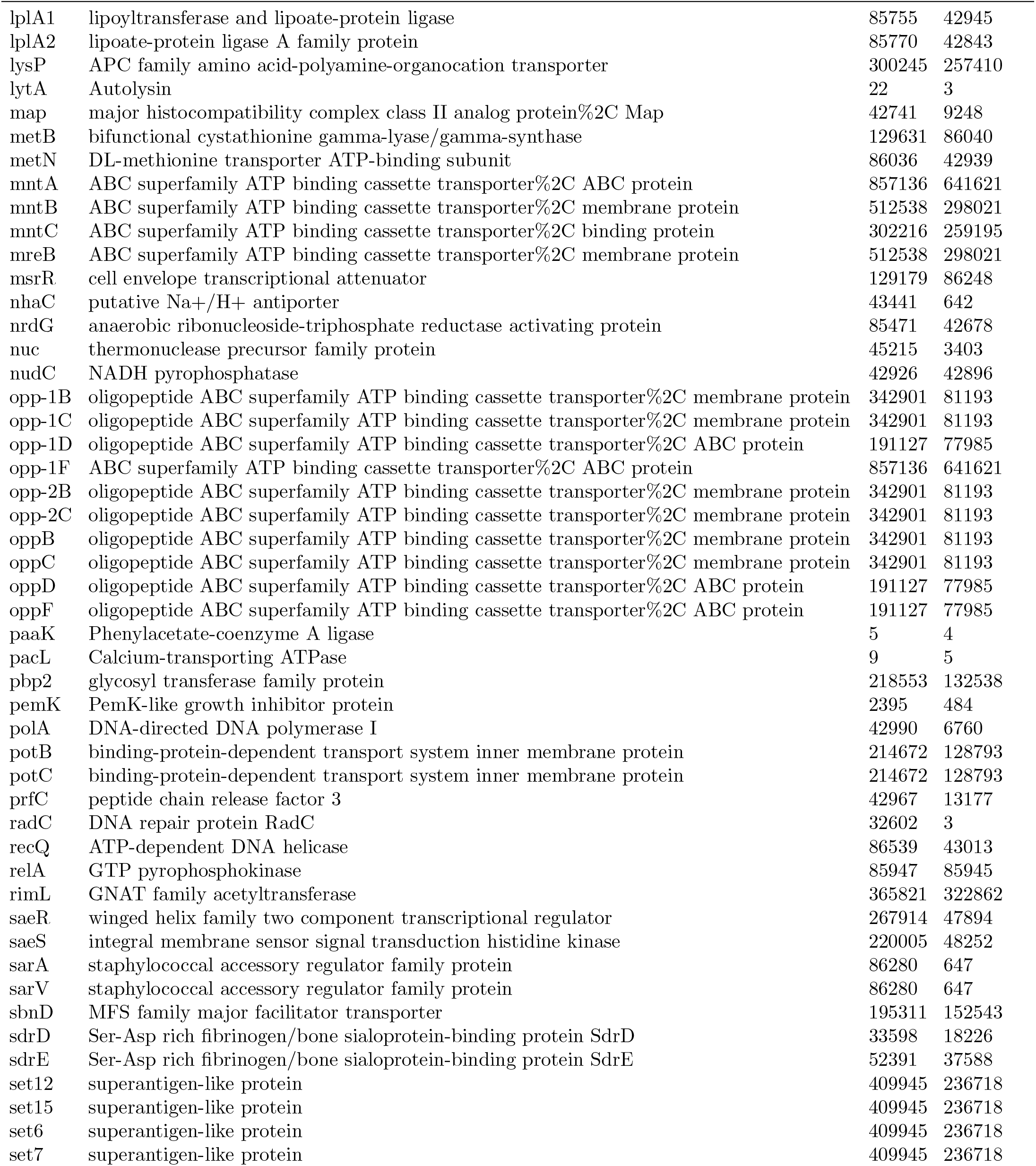
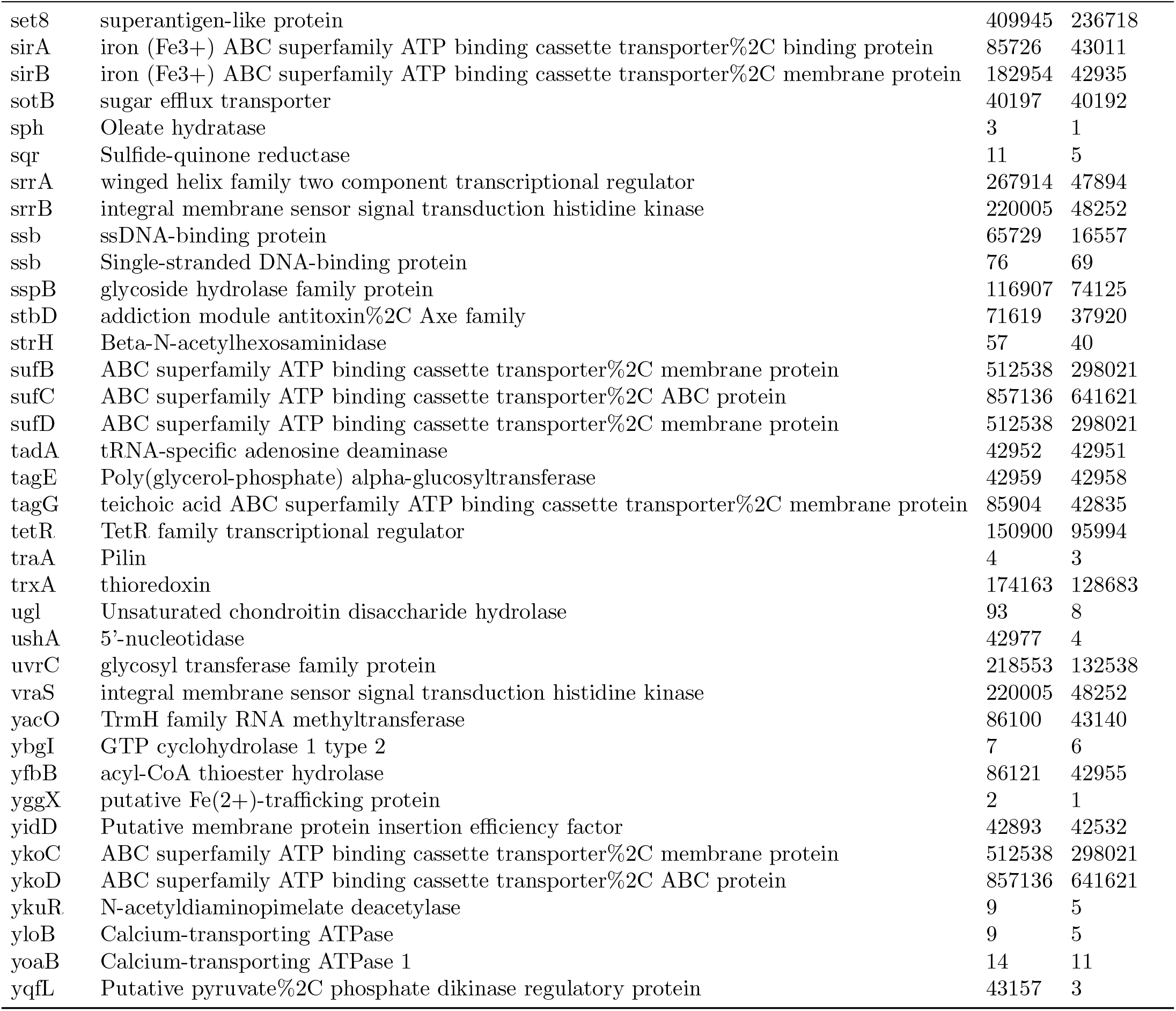
All named genes whose products also corresponded to unnamed genes in Staphopia. Note that the “%2C”s are present in the database text.

## H Coevolution score does not systematically depend on the choice of distance cutoff

To study how the choice of distance cutoff affected the coevolution score, we computed the coevolution score for each clonal complex for each gene pair across a range of distance cutoffs, chosen such that for each clonal complex separately, the number of close pairs used ranged from 1000 to 10,000 in intervals of 1000. We chose the distance cutoff that results from using 5000 close pairs for Table S1.

Figure S5 displays the scores as a function of distance cutoff for clonal complex CC15. The scores for individual gene interactions are connected by lines. While specific rank order may change, in general, the large scores remain large and the small scores remain small across the range of distance cutoffs. The distance cutoff we have chosen may be conservative by this analysis.

**Figure S5:**
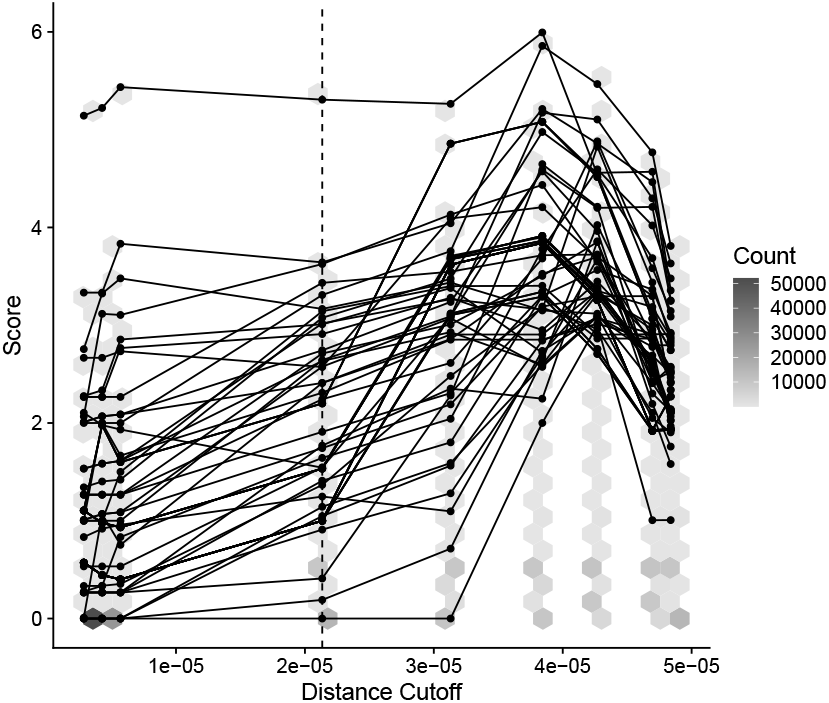
The choice of the distance cutoff does not systematically affect the coevolution score, and the highest scores remain high across a wide range of cutoffs. For gene pairs whose maximum score across these cutoffs is at least 3, points that correspond to the same gene pair are connected by lines. All other gene pairs are represented in the hexagonal heatmap. The dashed line depicts the cutoff for 5000 close pairs used. These gene pairs are from clonal complex CC15.

## I Full-dataset analysis

Because scales of divergence vary dramatically between clonal complexes (Figure S2), simply choosing a single distance cutoff using all nonredundant samples from the dataset will bias the set of chosen samples towards dramatic overrepresentation of some clonal complexes but not others. Choosing a larger distance cutoff leads to a more representative sample (Figure S6).

On the other hand, choosing a larger distance threshold results in an enormous number of close pairs that render the procedure to compute coevolution scores computationally intensive. To combat this problem, we chose a compromise distance threshold of 0.0005 (vertical line in Figure S6). In choosing this distance threshold, we did not consider CC5, CC8, or CC22; these clonal complexes had many samples that were identical in the core sequence. For the other clonal complexes, we found that there were approximately 18 million close pairs at this distance cutoff, resulting in approximately 1.8 million per clonal complex. For CC5, CC8, and CC22, we randomly sampled 1.8 million close pairs each from the set of close pairs that were below the distance cutoff, resulting in a total of approximately 23.4 million close pairs. From this total set of close pairs, we randomly sampled 40,000 to serve as our set of close pairs for the full dataset analysis.

**Figure S6:**
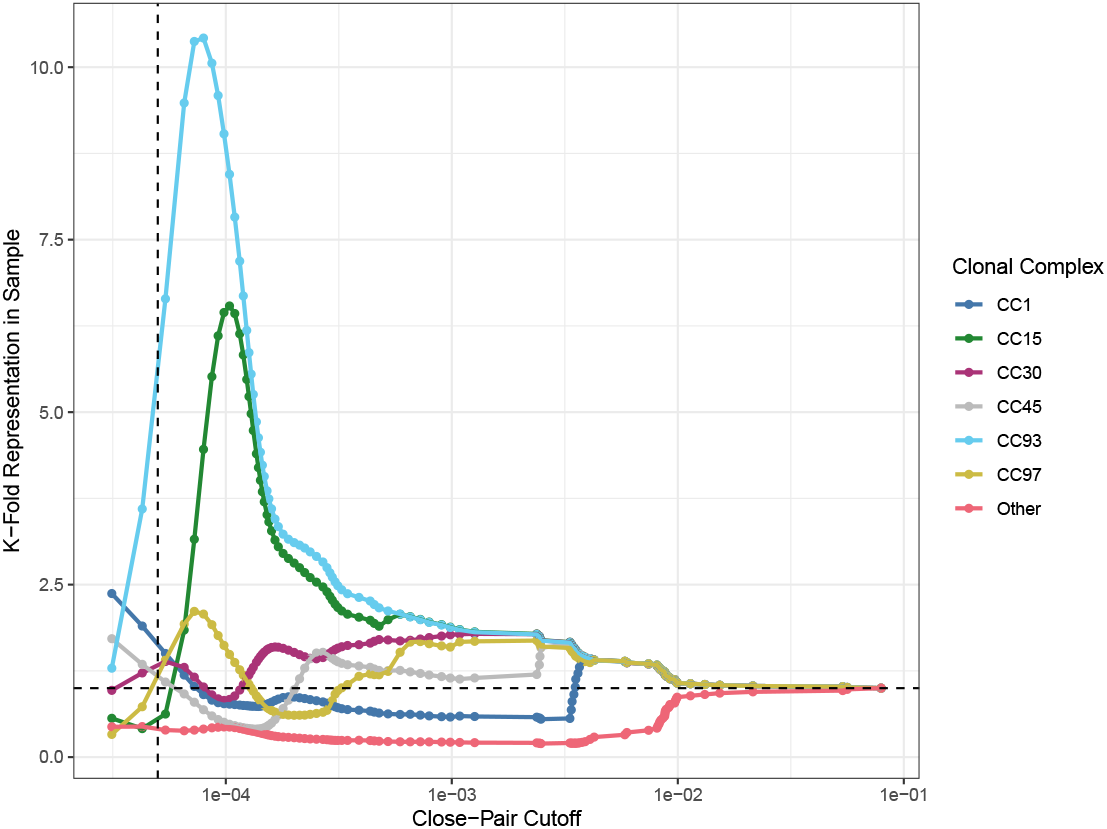
Choosing a core-genome distance cutoff across the whole Staphopia dataset overrepresents some clonal complexes and underrepresents others. The y-axis represents the ratio of the fraction of samples in each clonal complex below the corresponding distance cutoff to the fraction of samples in each clonal complex in the whole dataset. Excluded from this set of clonal complexes are CC5, CC8, and CC22, which have substantial numbers of samples that are identical in the core genome. The vertical line represents the distance cutoff chosen for the full dataset analysis.

